# A two-hit epistasis model prevents core genome disharmony in recombining bacteria

**DOI:** 10.1101/2021.03.15.435406

**Authors:** Aidan J. Taylor, Koji Yahara, Ben Pascoe, Leonardos Mageiros, Evangelos Mourkas, Jessica K Calland, Santeri Puranen, Matthew D. Hitchings, Keith A. Jolley, Carolin M. Kobras, Sion Bayliss, Nicola J. Williams, Arnoud H. M. van Vliet, Julian Parkhill, Martin C. J. Maiden, Jukka Corander, Laurence D Hurst, Daniel Falush, Paul Keim, Xavier Didelot, David J. Kelly, Samuel K. Sheppard

**Author notes:** Corresponding author(s): David J. Kelly,; Samuel K. Sheppard.

## Abstract

**Significance Statement:** Genetic exchange among bacteria shapes the microbial world. From the acquisition of antimicrobial resistance genes to fundamental questions about the nature of bacterial species, this powerful evolutionary force has preoccupied scientists for decades. However, the mixing of genes between species rests on a paradox. On one hand, promoting adaptation by conferring novel functionality, on the other potentially introducing disharmonious gene combinations (negative epistasis) that will be selected against. Taking an interdisciplinary approach to analyse natural populations of the enteric bacteria *Campylobacter*, an ideal example of long-range admixture, we demonstrate that genes can independently transfer across species boundaries and rejoin in epistasis in a recipient genome. This challenges conventional ideas and highlights the possibility of single step evolution by saltation.

**Abstract:** Recombination of short DNA fragments via horizontal gene transfer (HGT) can both introduce beneficial alleles and create genomic disharmony through negative epistasis. For non-core (accessory) genes, the negative epistatic cost is likely to be minimal because the incoming genes have not co-evolved with the recipient genome. By contrast, for the core genome, interspecific recombination is expected to be rare because disruptive allelic replacement is likely to introduce negative epistasis. Why then is homologous recombination common in the core of bacterial genomes? To understand this enigma we take advantage of an exceptional model system, the common enteric pathogens *Campylobacter jejuni* and *Campylobacter coli*, that are known for very high magnitude interspecies gene flow in the core genome. As expected, HGT does indeed disrupt co-adapted allele pairings (negative epistasis). However, multiple HGT events enable recovery of the genome’s co-adaption between introgressing alleles, even in core metabolism genes (e.g., formate dehydrogenase). These findings demonstrate that, even for complex traits, genetic coalitions can be decoupled, transferred and independently reinstated in a new genetic background – facilitating transition between fitness peaks. In this example, the two-step recombinational process is associated with *C. coli* that are adapted to the agricultural niche.

## Introduction

Mutation is the engine of genetic novelty, but for most bacteria adaptation also involves the acquisition of DNA from other strains and species through horizontal gene transfer (HGT) [1]. In some wel documented cases, a single nucleotide substitution or acquisition of a small number of genes, can prompt new evolutionary trajectories with striking outcomes such as the evolution of virulent or antibiotic resistant strains [2]. With such dynamic genomic architecture, it may be tempting (and possibly useful [3]) to consider genes as independent units that ‘plug and play’ innovation into recipient genomes. This is clearly an oversimplification. In fact, genes within genomes are highly interactive wherein the effect of one allele depends on another (epistasis). Therefore, it is likely that some horizontally introduced changes will disrupt gene networks and be costly to the original coadapted genetic background, especially for complex phenotypes involving multiple genes and more distant taxonomic relationships.

Understanding how epistasis influences the evolution of phenotype diversity has preoccupied researchers since the origin of population genetics [4–10], with much emphasis placed upon the relative amounts of recombination and epistatic effect sizes [11, 12]. In sexual populations, such as outbreeding metazoans, genetic variation is shuffled at each generation. This the trend towards randomization between alleles (linkage equilibrium), means it is unlikely that multiple distinct epistatic allele combinations will be maintained in the same population [8]. In bacteria, however, rapid clonal reproduction allows multiple genomically distant beneficial allele combinations to rise to high frequency or even fixed in a single population. For example, in common enteric bacteria such as *Escherichia coli, Salmonella enterica* and *Campylobacter jejuni,* the doubling time in the wild has been estimated at around 24 hours or less [13, 14]. Therefore, though HGT occurs in these organisms [15], even in highly recombinogenic *C. jejuni* [13, 16], there will likely be many millions of bacterial generations between recombination events at a given locus. This allows mutations that are beneficial only in specific genetic backgrounds to establish in a single population and linkage disequilibrium to form between different epistatic pairs [17].

In a coadapted genomic landscape, recombination is expected to have two antagonistic effects. On one hand it could be beneficial, promoting adaptation by conferring novel functionality on the recipient genome [18–22] and though specialization reduce the competition between clones (clonal interference). On the other it could be detrimental, introducing disharmonious allele-combinations that will be discriminated against by selection [19]. The balance of these two effects is likely to be different for core and non-core HGT events. Non-core events, such as introduced accessory antibiotic resistance genes, can be immediately beneficial and, as they do not replace recipient sequence, need not break established epistatic interactions. By contrast, HGT replacing one allele for another (homologous recombination) in part of the core genome is likely to disrupt highly evolved co-adapted networks. For these reasons, negative epistatic interactions between core genes with different evolutionary histories have been proposed as a barrier to recombination [4–10, 23], particularly between species. However, interspecies recombination is common in bacterial core genomes [18, 24]. How then can a core genome, that is expected to build up extensive co-adapted epistatic networks, be so accepting of HGT events?

The common animal gut bacterium *Campylobacter,* which is among the most prolific causes of human bacterial gastroenteritis worldwide [25], provides an exceptional model system to study the impact of HGT on the core genome for several reasons. First, introgression between the two most important pathogenic species, *C. jejuni* and *C. coli,* has led to the evolution of a globally distributed ‘hybrid’ *C. coli* lineage [26] that is responsible for almost all livestock and human infections with this species. The hybrids occupy a niche that was not available to the parental subtypes. Second, up to 23% of the core genome of one common *C. coli* subtype has been recently introgressed from *C. jejuni* [27], potentially disrupting epistatic interactions. Finally, because *C. jejuni* and *C. coli* have undergone an extended period of independent evolution with only 85% average nucleotide identity, recombined sequence is conspicuous in the genome.

Here, we investigate the disruptive effect of HGT on co-adapted bacterial core genomes by examining covarying allele pairings and imported DNA sequence. Even though recombined fragments enter the genome one-by-one, we find that most covariation in the core genome is between sites where both alleles were imported. Having confirmed that allelic covariation is indicative of epistasis, with laboratory mutagenesis and complementation assays, we conclude that independent disruptive recombination events occur and persist until a second event restores the functional link in a new genetic background. This process resembles the two-hit cancer model where an initial mutation is actuated by a second [48]. Both bacterial and cancer cell lines are asexual clones and a two-hit model for core genome HGT is consistent with a more general theory in which negative effects from an initial event (MGT or mutation) need not preclude adaptive evolution. As in cancer, the recombinant genotypes are under strong selection and occupy a unique agricultural niche.

## Results

### *Campylobacter* populations are highly structured with intermediate sequence clusters

Analysis of every automatically annotated gene from every genome, revealed a core genome of 631 gene orthologues in all *Campylobacter* isolates in this study. There were 1287 genes common to all *C. jejuni* isolates and 895, 1021 and 1272 common to *C. coli* clades 1, 2 and 3, respectively. Consistent with previous studies [27], neighbour-joining and ClonalFrameML trees based on genes within concatenated core genome alignments revealed population structure in which *C. jejuni* and *C. coli* clade 2 and 3 isolates each formed discrete clusters (Figure 1A, Figure S1). Isolates designated as *C. coli* clade 1 were found in three clusters on the phylogeny: unintrogressed ancestral strains, and the ST-828 and ST-1150 clonal complexes (CCs) which account for the great majority of strains found in agriculture and human disease [27]. Pan-genome analysis quantified the increase in unique gene discovery as the number of sampled genomes increased (Figure S2A). For all phylogenetic clusters there was evidence of an open pan-genome with a trend towards continued rapid gene discovery within clusters containing fewer isolates. There was considerable accessory genome variation between species and clades, potentially associated with important adaptive traits (Figure S3) and there was evidence that the average number of genes per genome was greater in *C. coli* CC-828 isolates than in *C. jejuni* (Figure S2B).

**Figure 1.**
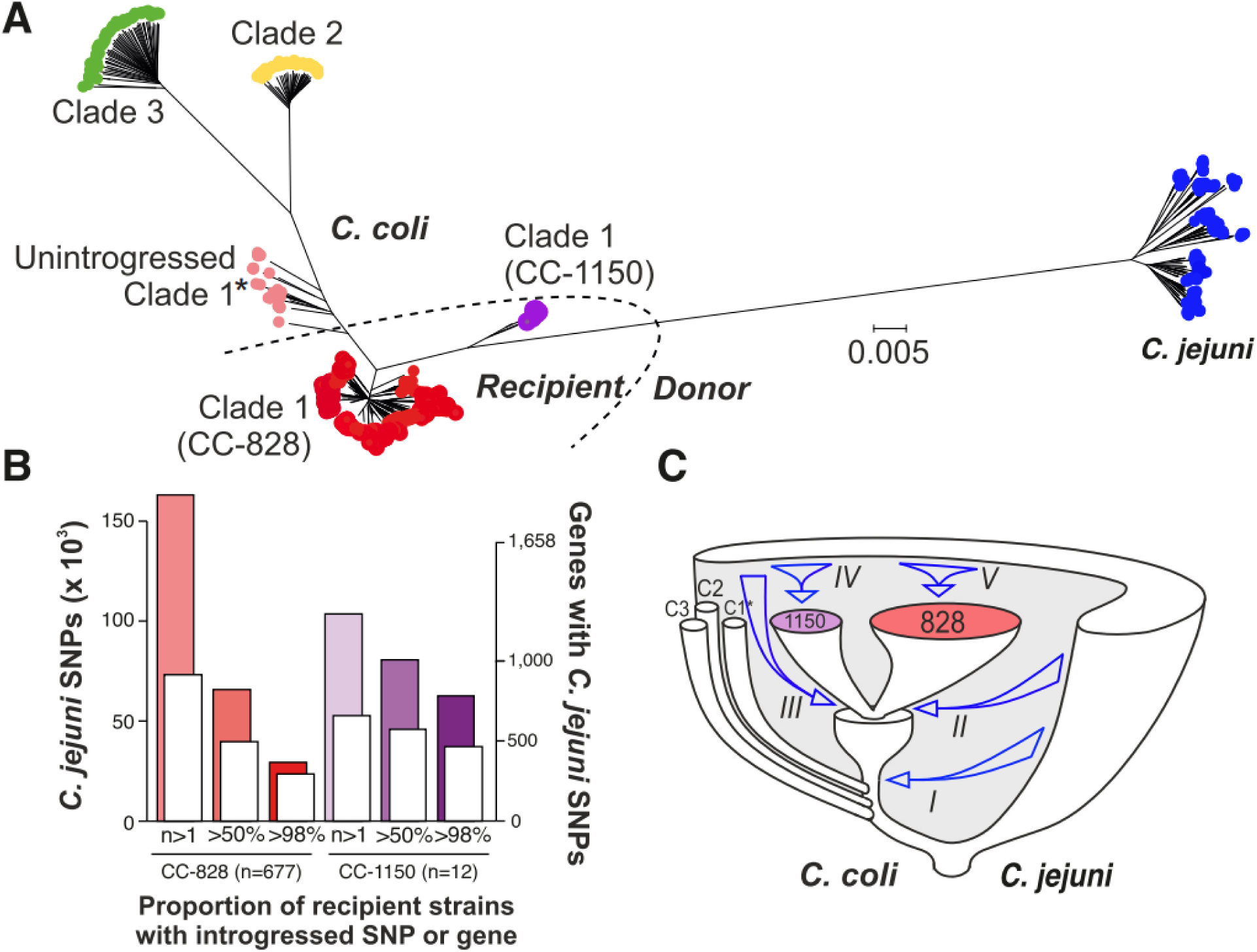
Genome-wide introgression from *C. jejuni* to *C. coli*. (A) Phylogenetic tree reconstructed using neighbour-joining on a whole-genome alignment of 973 *C. jejuni* and *C. coli* isolates. Introgressed *C. coli* clades are represented with red (CC-828, n=677) and purple (CC- 1150, n=12) circles, unintrogressed clade 1* (n=35) is shown in pink, clade 2 (n=45) in yellow and clade 3 (n=58) in green. A set of 146 *C. jejuni* genomes belonging to 30 clonal complexes (4 to 5 isolates per ST) are shown in blue. Recipient and donor populations, used to infer introgression in chromosome painting analysis, are indicated. The scale bar represents the number of substitutions per site. (B) Summary of introgressed *C. jejuni* SNPs in *C. coli* CC-828 (n=677, red) and CC-1150 (n=12, purple) genomes using ChromoPainter; the number of introgressed core SNPs (coloured histograms; left y-axis) and core genes (white histograms; right y-axis) for a range of recipient strains proportions (at least 1, more than 50% and more than 98%) is shown. (C) Diagram of *Campylobacter* species and clade (C1*, C2, C3) divergence with arrows indicating sequential introgression from *C. jejuni* into *C. coli* (I) clade 1, (II, III) CC-828 and CC- 1150, (IV, V) subsequent clonal expansion and ongoing introgression.

### There is substantial introgression within the *C. coli* core genome

The large genetic distances among *C. coli* clade 1 isolates (Figure 1A), have been shown to be a consequence of the import of DNA from *C. jejuni* rather than accumulation of mutations during a prolonged period of separate evolution [26]. Using chromosome painting to infer the co-ancestry of core-genome haplotype data from CC-828 and CC-1150 isolates gave a detailed representation of the recombination-derived segments from each *C. jejuni* donor group and clonal complex to each recipient individual (Figure S4). The majority of introgressed SNPs were rare, occurring in fewer than 50 recipient genomes (Figure 1B). However, a large proportion of the introgressed *C. coli* clade 1 genomes contained DNA of *C. jejuni* ancestry in >98% of recipient isolates. Consistent with previous estimates [27], these regions where introgressed DNA was largely fixed within the *C. coli* population, occurred across the genome and comprised up to 15% and 28% of the CC-828 and CC-1150 isolate genomes, respectively (Figure 1B). After considering potential donor groups, we find that the majority of introgressed DNA in *C. coli* involves genes that are present in multiple *C. jejuni* lineages (core genes) (Figure S5A).

Having identified *C. jejuni* ancestry within *C. coli* genomes, we investigated the sequence of events responsible for introgression. Most introgressed SNPs are found at low frequency in both CC-828 and CC-1150 clonal complexes. However, there was evidence of SNPs that are introgressed in both complexes as well as high frequency lineage specific introgression in both CC-828 and CC- 1150 (Figure S5B). Specifically, 25% of the *C. jejuni* DNA found in >98% of CC-828 isolates was also found in CC-1150 (Figure S5C), implying that this genetic material was imported by the common ancestor(s) of both complexes. After the divergence of these two complexes, introgression continued with nearly 75% of *C. jejuni* DNA present in one recipient clonal complex and not the other. This stepwise model is consistent with an evolutionary history in which there was a period of progressive species and clade divergence reaching approximately 12% at the nucleotide level between *C. jejuni* and *C. coli* and around 4% between the three *C. coli* lineages. More recently, changes to the patterns of gene flow led one *C. coli* clade 1 lineage to import substantial quantities of *C. jejuni* DNA, and further lineage-specific introgression gave rise to two clonal complexes (CC-828 and CC-1150) that continued to accumulate *C. jejuni* DNA, independently creating the population structure observed today (Figure 1C).

The high magnitude introgression into *C. coli* clade 1 isolates has introduced thousands of nucleotide changes to the core genome. However, divergence in bacteria may be uneven across the genome for at least two reasons. First, recombination is more likely to occur in regions where donor and recipient genomes have high nucleotide similarity [24, 28, 29]. Second, ‘fragmented speciation’ [30], in which gene flow varies in different parts of the genome, such as regions responsible for adaptive divergence, can result in phylogenetic incongruence among genes. Consistent with previous estimates [27], we found that the three *C. coli* clades had similar high divergence with *C. jejuni* across the genome, ranging from 68% to 98% nucleotide identity for individual genes (Figure S5D), implying a period of divergence with low levels of gene flow. We found no evidence that high genetic differentiation between the species prevented recombination. While there was some evidence that more recombination occurred in regions of low nucleotide divergence (between unintrogressed *C. coli* clade 1 and *C. jejuni*), introgression occurred across the genome at sites with a wide range of nucleotide identity (Figure S5D). This level of recombination has greatly increased overall genetic diversity across the genome in *C. coli* clade 1 and introduced changes that have potential functional significance.

### Much of the putative epistasis occurs between SNPs in introgressed genes

ClonalFrameML analysis revealed the importance of homologous recombination in generating sequence variation within the introgressed *C. coli*. Estimates of the relative frequency of recombination versus mutation (R/θ=0.43), mean recombination event length (δ=152bp) and average amount of polymorphism per site in recombined fragments (ν=0.07), imply that recombination has had an effect (*r/m*) 4.57 times higher than *de novo* mutation during the diversification of CC-828. This is consistent with previous analysis and confirmed recombination as the major driver of molecular evolution in *C. coli* [13, 27, 31]. The continuous time Markov chain model for the joint evolution of pairs of biallelic sites on a phylogenetic tree (Figure S6) was applied to investigate patterns of covariance for all pairs of sites >20kb apart (Figure S7). For most biallelic sites there were few branches on the tree where substitutions occurred, so that their evolution is compatible with separate evolution on the same clonal frame. However, 2874 covarying pairs evolved more frequently together than would be expected (*p-*value 10^-8^) if they had evolved independently based on the tree, and hence indicated patterns of putative epistasis.

Among them, the location of 2618 putative epistatic pairs of sites was compared to the inferred ancestry (unintrogressed *C. coli* or *C. jejuni*) of sequence across the genome of CC-828 and CC- 1150 *C. coli* strains (Figure 2C, Data S1). Putative epistatic pairs in the recipient introgressed genome represent a combination of the haplotypes in the ancestral populations (Figure 2A). For each epistatic pair, the major and minor haplotype were defined, from these possible combinations, if there was haplotype polymorphism between *C. jejuni* and *C. coli.* This allowed quantification of the number of covarying sites that occurred between an ancestral *C. coli* (unintrogressed) and an introgressed *C. jejuni* allele (*C. coli – C. jejuni*), two introgressed alleles (*C. jejuni – C. jejuni*), and sites that do not segregate by species. Strikingly, the breakdown of the major and minor haplotype combinations among the 2618 epistatic pairs (Figure 2B, Data S2) shows the major haplotype for 83.5% of putative epistatic SNP pairs was *C. jejuni* indicating that both co-varying sites had *C. jejuni* ancestry, consistent with epistasis between introgressed ancestral *C. jejuni* sequence at divergent genomic positions. Investigation of the genes containing co-varying sites revealed that 2187 SNP pairs were in 16 genes with just five genes accounting for 99.1% of them (Figure 3A&B, Data S3, Figure S8).

**Figure 2.**
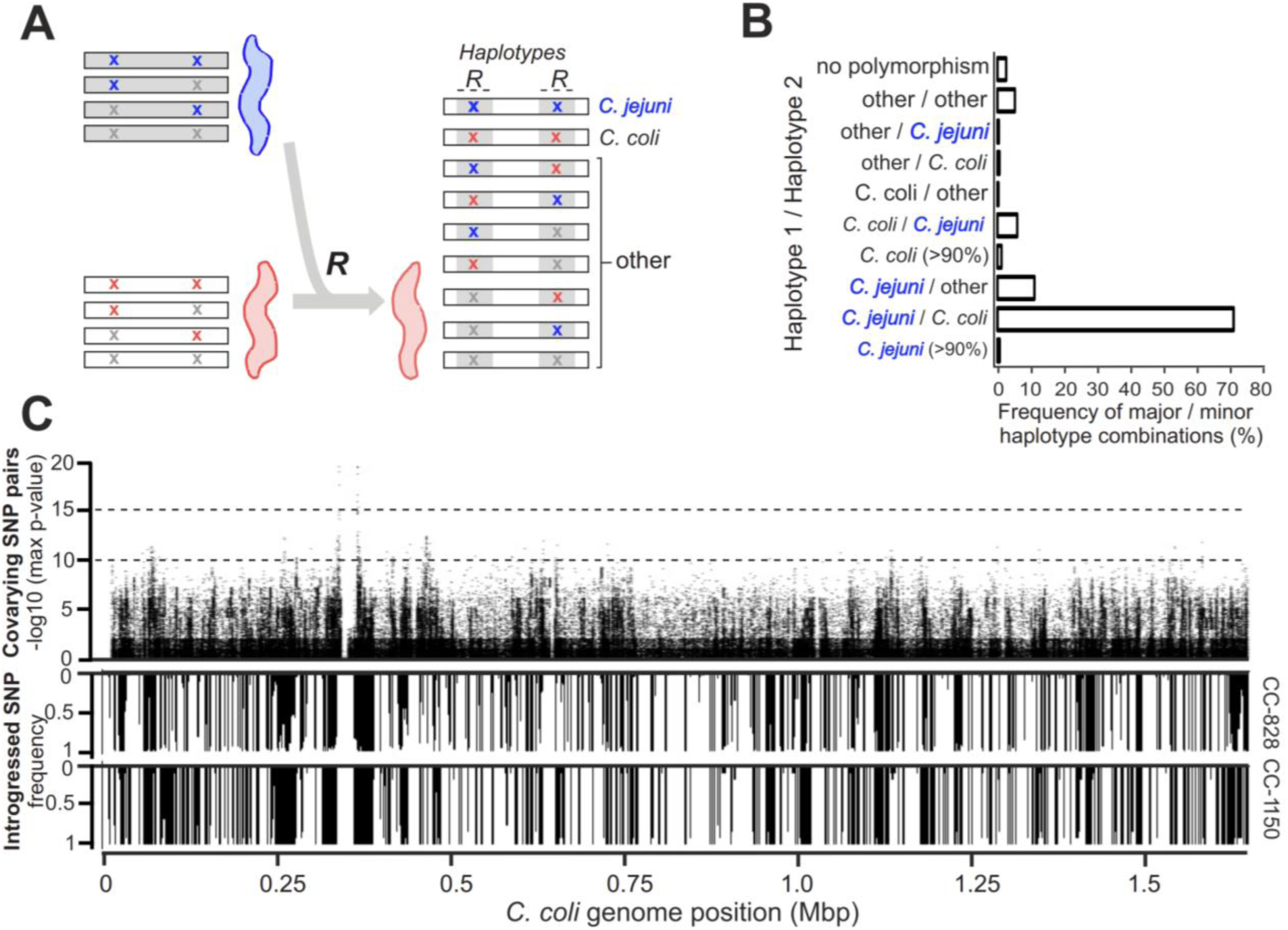
Covariation in introgressed *C. coli* genomes. (A) Possible SNP pairs in ancestral donor *C. jejuni* and recipient *C. coli* genomes and potential recombinant (***R***) haplotype pairs in introgressed *C. coli.* Possible genome haplotypes are designated with *C. jejuni* fixed SNPs (blue X), other *C. jejuni* SNP (gray X), *C. coli* fixed SNP (red X), and other *C. coli* SNPs (gray X). (B) The frequency of major and minor haplotype combinations among the 2578 pairs of covarying SNPs in the 689 *C. coli* clade-1 recipient genomes, revealing that 83.5% of long-range covariation was between introgressed *C. jejuni* sites. (C) Miami plot of each polymorphic site showing the maximum *p-*value for covarying biallelic pairs (>20kb apart) and the frequency of introgression in CC-828 and CC-1150.

**Figure 3.**
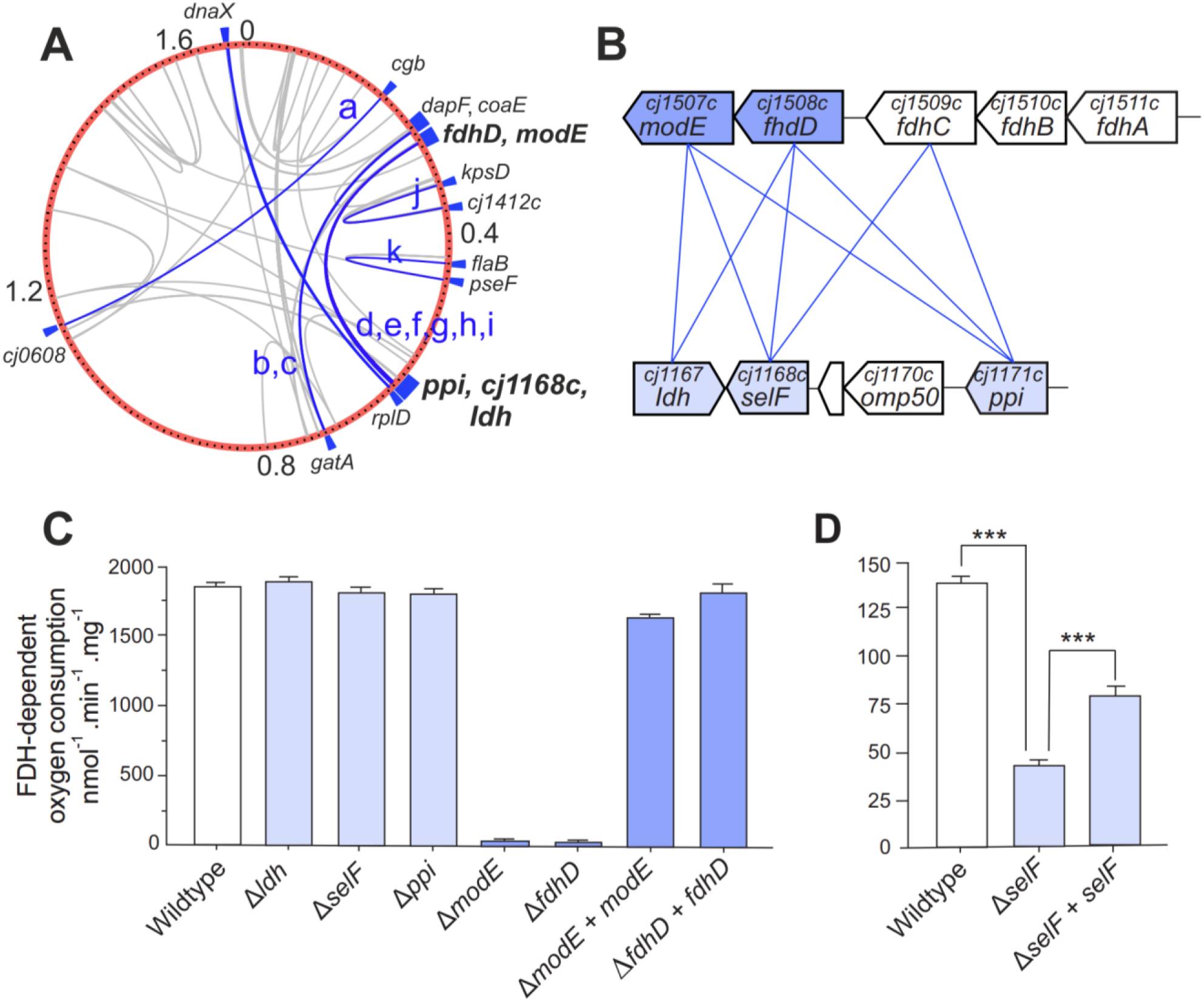
Genomic context and physiological roles of introgressed epistatically linked genes involved in FDH biogenesis and activity. (A) The position of putative epistatic sites mapped on the *C. coli* CVM29710 reference for covarying *C. jejuni-C. jejuni* SNPs (blue) in 16 gene pairs (a to l), and other haplotype combinations (grey). (B) Genome organization of the most common co varying introgressed *C. jejuni* – *C. jejuni* gene pairs linked to FDH. (C-D) FDH activity of whole cells determined by oxygen consumption rates in a Clark-type electrode (nmol oxygen consumed per minute per mg of total protein) for (C) cells grown in rich media (excess selenium), (D) cells grown in minimal media with 0.5 nM sodium selenite.

### Genomic context and physiological role of epistatically linked genes

The five genes accounting for the majority of epistatic interactions (*cj1167*, *cj1168c*, *cj1171c*/*ppi*, *cj1507c*/*modE* and *cj1508c*/*fdhD*) were investigated for their physiological role in *C. jejuni*. FdhD and ModE are proteins involved in the biogenesis of formate dehydrogenase (FDH). The FDH complex (FdhABC) oxidises formate to bicarbonate to generate electrons that fuel cellular respiration. Formate is an abundant electron donor produced by host microbiota and an important energy source for *Campylobacter in vivo* [32, 33]. The remaining three genes, *cj1167* (annotated incorrectly as *ldh*, lactate dehydrogenase), *cj1168c* and *cj1171c* (*ppi*) are also grouped together in the genome, where *cj1167* and *cj1168c* are adjacent but with the open reading frames (ORFs) convergent and overlapping, while *ppi* is upstream, separated by two non-epistatically linked genes (*cj1169c* and *cj1170c*, Figure 3B). Considering the genomic arrangement, it is therefore clear that the putative epistatic links uncovered in this study essentially occur across two loci in the genome (*fdhD*/*modE* and *cj1167*/*cj1168c*/*ppi*), with each of the latter three genes linked with both *fdhD* and *modE* (Figure 3B). Given the known function of *fdhD* and *modE* in biogenesis of the FDH complex, we hypothesised that *cj1167*/*cj1168c*/*ppi* might also have some role in FDH biogenesis or activity in order to form a functional epistatic connection. We therefore constructed deletion mutants to investigate the possible role of these genes in FDH activity.

Initially, each of the mutants and their parental wildtype (*C. jejuni* NCTC11168) were grown in rich media (Muller-Hinton broth) and their formate dependent oxygen consumption rates determined (Figure 3C). *cj1167, cj1168c* and *ppi* mutants demonstrated wildtype levels of FDH activity, while activity in both *fdhD* and *modE* mutants was abolished. To confirm that the phenotype of the *fdhD* and *modE* mutants was not due to a polar effect on the surrounding *fdh* locus, these mutants were genetically complemented by reintroduction of a second copy of the wildtype gene into the rRNA locus, which restored near wildtype levels of FDH activity in both cases (Figure 3C).

As neither *cj1167, cj1168c* or *ppi* mutants showed altered FDH activity in cells grown in rich media, we considered that their function may be related to an FDH-specific nutrient requirement as would likely be found *in vivo*. Since the formate oxidising subunit of FDH, FdhA, specifically requires a molybdo or tungsto-pterin (Mo/W) cofactor and a selenocysteine (SeC) residue for catalysis [34], Mo, W or Se supply presented possible targets. *cj1168c* encodes a DedA family integral membrane protein of unknown function. DedA proteins are solute transporters widespread in bacteria but are mostly uncharacterized [35]. However, a homologue of *cj1168c* in the heavy metal specialist beta-proteobacterium *Cupriavidus metallidurans* has been shown to be involved in selenite (SeO_3_^2-^) uptake [36]. We therefore speculated that Cj1168 could be a selenium oxyanion transporter that supplies Se for SeC biosynthesis. To test this, FDH activities were measured in *cj1168c* mutant and parental wildtype strains grown in minimal media with limiting concentrations of selenite or selenate (SeO_4_^2-^) (Figure S10). The data in Figure 3D shows that the *cj1168c* mutant displayed significantly reduced FDH activity after growth with selenite in the low nM range, and this phenotype was partially restored by genetic complementation. We therefore designated *cj1168c* as *selF* (<u>se</u>lenium transporter for formate dehydrogenase). However, although this phenotype does suggest that SelF is a selenium importer, another unrelated selenium transporter, FdhT (Cj1500), has previously been documented in *C. jejuni* [37], which is not epistatic with *fdhD* or *modE*. In contrast to this previous report, we found considerable residual FDH activity still remained in an *fdhT* deletion mutant, which was fully restored to wildtype levels by complementation (Figure S9).

Finally, we tested whether the residual FDH activity in our *fdhT* mutant was due to selenium uptake by SelF. An *fdhT selF* double mutant was generated and assayed for FDH activity after growth in minimal media containing limiting concentrations of selenite or selenate (Figure S9). The *fdhT selF* double mutant demonstrated a significant additional reduction in FDH activity over the *fdhT* single mutant, a phenotype that was partially restored by complementing the double mutant with *selF*. Complementation of the double mutant with *fdhT* returned FDH activity to near wildtype levels (Figure S9). Taken together, our data suggests that both FdhT and SelF facilitate selenium acquisition in *C. jejuni*, possibly representing low and high affinity transporters, respectively (Figure S11).

## Discussion

To address the apparent contradiction of frequent core genome recombination in a co-adapted genomic background we focused on *Campylobacter,* in which interspecies recombination is well documented [26, 27, 38]. As in other studies, we found that a large proportion (15-28%) of the *C. coli* core genome originated in *C. jejuni* despite the genetic distance (∼85% nucleotide identity) between the species. Investigating the likely disruptive impact this would have on coadapted epistatic gene networks, we quantified the frequency of *C. jejuni* – *C. coli* (and *vice versa*) and *C. jejuni* – *C. jejuni* covarying allele pairs in introgressed *C. coli*. Where recombination is minimally disruptive there would be more *C. jejuni* – *C. coli* than *C. jejuni* – *C. jejuni*. However, consistent with selection against disharmonious gene combinations we found that *C. jejuni* – *C. jejuni* allele pairs constituted >83% of covarying introgressed haplotypes. It is possible that in some cases both sites were introgressed in a single large recombination event [39–41] but in *Campylobacter* LD for pairs of sites decreases with distance, and is approximately constant after 20kbp [42] consistent with the independent acquisition of alleles that are >20kb apart. It follows, therefore, that the first introgression event was not fatal to the recipient genome. Acquisition of the second member of the pair then potentially restored the function of the integrated *C. jejuni* – *C. jejuni* coevolutionary unit.

Statistical genetic analysis confirmed covariation, but to confirm coadaptation (*sensu stricto)* it was necessary to confirm epistasis in the laboratory. The most common co-varying *C. jejuni* – *C. jejuni* gene pairs were linked to FDH, a key enzyme allowing the utilisation of formate as an electron donor *in vivo* [32, 33]. FdhD and ModE were shown to be essential for FDH activity.

While FdhD is a known sulfur-transferase required for cofactor insertion into FdhA [43] (Figure S11), the abolished FDH activity in a *modE* mutant indicates functions for ModE in FDH biogenesis, beyond the known role as a transcriptional repressor [44, 45]. In contrast, *cj1167, self* and *ppi* mutants did not show reduced FDH activity. This apparently contradicted the epistasis hypothesis until we considered the functional links between *fdhD*/*modE* and *cj1167*/*selF*/*ppi.* This revealed that a *selF* mutant strain had significantly reduced FDH activity under conditions of selenite limitation (Figure 3D). This phenotype is consistent with SelF being a Se oxyanion transporter and functionally links SelF with FdhD/ModE. We suggest that *selF* rather than *fdhT* is epistatic because SelF confers an additional benefit (SeC biosynthesis, essential for FDH activity) under conditions of selenium limitation, for example as may be found in the host gut (Figure S11).

Direct functional association with FDH activity was more difficult to explain for *cj1167* and *ppi. cj1167* is currently incorrectly annotated as lactate dehydrogenase (Ldh) and it’s actual function is unknown [46]. While we found no evidence for a functional connection between Cj1167 and FDH activity, the overlapping convergent gene arrangement of *cj1167* and *cj1168c* (*selF*) suggests a transcriptional architecture that might form similar epistatic dependencies even if Cj1167 is not required for FDH activity. Finally, the *cj1171c* (*ppi*) deletion mutant showed no growth defect or reduction in FDH activity. *ppi* encodes a cytoplasmic peptidyl-prolyl *cis-trans* isomerase and PPIases are general protein folding catalysts that often have pleiotropic and redundant functions [47]. Therefore, it is possible that if Cj1171 does help promote the folding of FdhD or ModE but analysis of a simple deletion mutant may not reveal this if another PPIase can substitute in that genetic background.

Understanding the functional significance of core genome recombination has considerable potential to explain the evolution of complex phenotypes. In *Campylobacter,* our results suggest that an ancestral *C. coli* lineage colonized a new niche and surviving lineages (CC-828 and CC- 1150) gained access to *C. jejuni* DNA (Figure 4A&B). In this case, introgressed *C. coli* acquired multiple genes allowing them to utilize the high levels of formate in the host gut as an energy source. Specifically, the *fdh* and *ModE,* essential for FDH biogenesis, as well as *selF* that ensures sufficient selenium for the enzyme to function. This example shows how as the adaptive landscape of the genome changed, potentially decoupling epistatic interactions that were previously selected, new gene combinations can be introduced by homologous recombination and tested in the *C. coli* genetic background. These new recombinant genotypes are frequently isolated in the agricultural setting and from clinical cases caused by agricultural products, arguing for their enhanced fitness in this niche.

**Figure 4.**
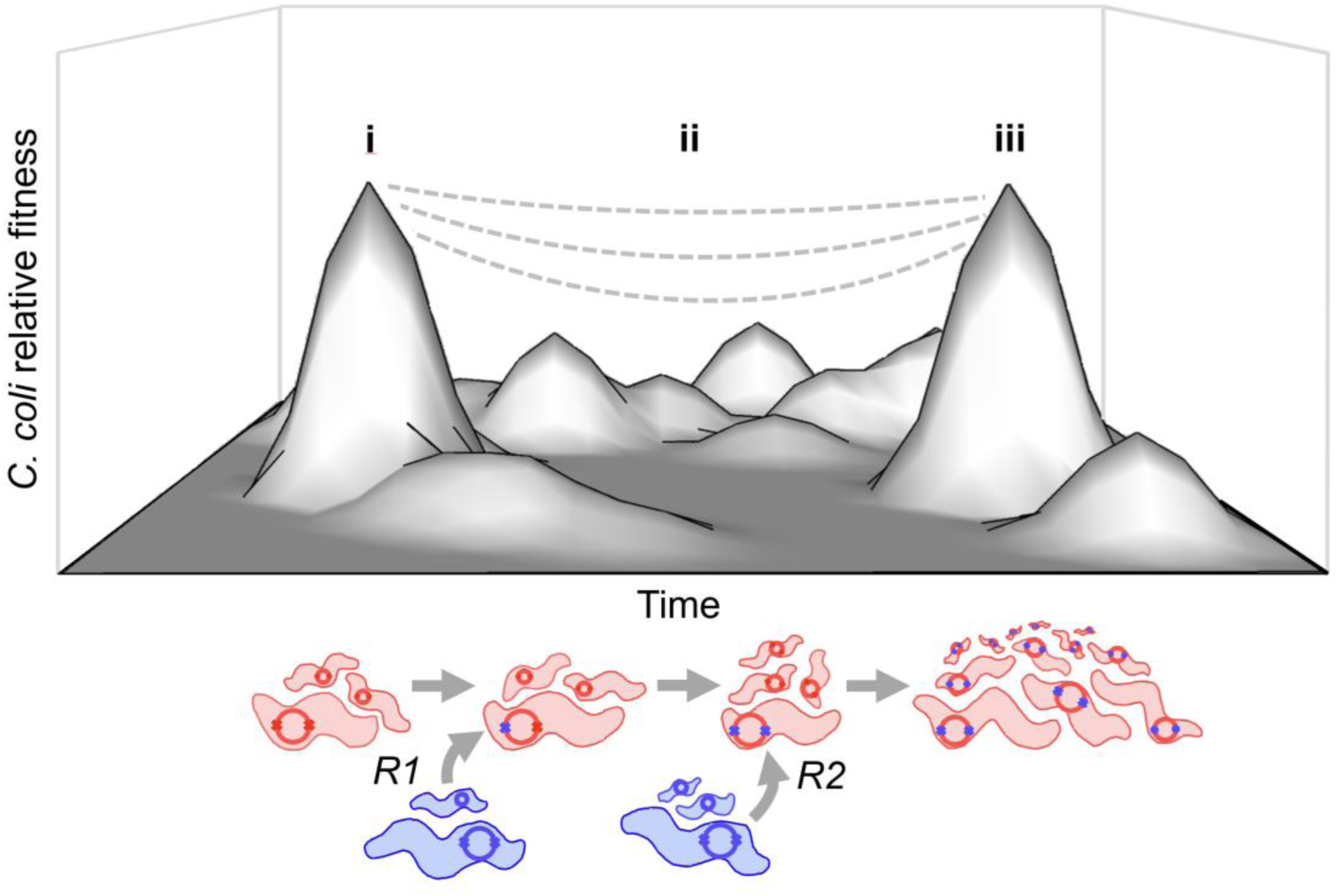
A two-hit model for core genome recombination in natural *C. coli* populations. Fitness landscape upon which (i) unintrogressed *C. coli* (red) and *C. jejuni* (blue) co-exist. Their genomes (internal circles) harbour haplotype pairs (x-x) that segregate by species. (ii) Horizontal gene transfer, HGT, occurs (*R1*) disrupting covarying genetic elements and reducing the relative fitness of introgressed *C. coli* to varying degrees, few strains retain mixed *C. coli - C. jejuni* haplotypes. (iii) HGT continues (*R2*) and, where recombined mixed haplotypes survived, ancestral *C. jejuni* haplotype pairs are reinstated in introgressed *C. coli –* some of which outcompete unintrogressed strains.

The notion that multiple events are necessary to achieve a phenotype with the first event potentially being deleterious (causing negative epistatsis) is strongly reminiscent of the two-hit cancer model in eukaryotes [48] in which the benefit to the tumour cell appears only after earlier non advantageous mutations. In bacteria, while the first hit (HGT event) in a core genome, is consistent with fitness costs associated with negative epistasis (breaking co-adapted gene networks), this can be rescued by a second HGT event. The second event, in effect restores a pre-existing co-adapted allele pair from the donor species to the recipient. This type of genetic rewiring may indeed be more common than previously thought [49–51] and, despite theoretical expectations, negative epistasis is not an absolute barrier to genome-wide recombination in structured bacterial populations. Multiple HGT events thus provide a solution to the classic evolutionary fitness peak jumping problem.

## Materials and Methods

### Isolates, genome sequencing and assembly

A total of 973 isolates were used in this study, 827 from *C. coli* and a selection of 146 from a diversity of *C. jejuni* clonal complexes (Data S4). Isolates were sampled mostly in the United Kingdom to maximise environmental and riparian reservoirs and thus the representation of genetic diversity in *C. coli*. Isolates were stored in a 20% (v/v) glycerol medium mix at -80°C and subcultured onto *Campylobacter* selective blood-free agar (mCCDA, CM0739, Oxoid). Plates were incubated at 42°C for 48 h under microaerobic conditions (5% CO2, 5% O2) generated using a CampyGen (CN0025, Oxoid) sachet in a sealed container. Subsequent phenotype assays were performed on Brucella agar (CM0271, Oxoid). Colonies were picked onto fresh plates and genomic DNA extraction was carried out using the QIAamp® DNA Mini Kit (QIAGEN; cat. number: 51306) according to the manufacturer’s instructions. DNA was eluted in 100–200 μl of the supplied buffer and stored at −20°C. DNA was quantified using a Nanodrop spectrophotometer and high-throughput genome sequencing was performed on a MiSeq (Illumina, San Diego, CA, USA), using the Nextera XT Library Preparation Kit with standard protocols involving fragmentation of 2 μg genomic DNA by acoustic shearing to enrich for 600 bp fragments, A-tailing, adapter ligation and an overlap extension PCR using the Illumina 3 primer set to introduce specific tag sequences between the sequencing and flow cell binding sites of the Illumina adapter.

DNA cleanup was carried out after each step to remove DNA < 150 bp using a 1:1 ratio of AMPure^®^ paramagnetic beads (Beckman Coulter, Inc., USA). Short read paired-end data was assembled using the *de novo* assembly algorithm, SPAdes (version 3.10.0) [52]. All novel genome sequences (n=475) generated for use in this study are available on NCBI BioProjects PRJNA689604 and PRJEB11972. These were augmented with 498 previously published genomes and accession numbers for all genomes can be found in Data S4 [16, 27, 31, 42, 53–57].

### Genome archiving, pan-genome content analyses and phylogenetic reconstruction

Contiguous genome sequence assemblies were individually archived on the web-based database platform PubMLST [58] and sequence type (ST) and clonal complex (CC) designation were assigned based upon the *C. jejuni* and *C. coli* multi-locus sequence typing scheme [59]. To examine the full pan-genome content of the dataset, a reference pan-genome list was assembled as previously described [60]. Briefly, genome assemblies from all 973 genomes in this study were automatically annotated using the RAST/SEED platform[61], the BLAST algorithm was used to determine whether coding sequences from this list were allelic variants of one another or ‘unique’ genes, with two alleles of the same gene being defined as sharing >70% sequence identity on >10% of the sequence length. The prevalence of each gene in the collection of 973 genomes was determined using BLAST with a positive hit in a genome being defined as a local alignment of the reference sequence with the genomic sequence of >70% identity on >50% of the length, as previously described [62]. The resulting matrix was analysed for differentiating core and accessory genome variation. Genes present in all genomes were concatenated to produce a core-genome alignment, used for subsequent phylogenetic reconstructions. Phylogenetic trees were reconstructed using an approximation of maximum-likelihood phylogenetics in FastTree2 [63]. This tree was used as an input for ClonalFrameML [64] to produce core genome phylogenies with branch lengths corrected for recombination.

### Inference of introgression

All 973 genomes were aligned to a full reference sequence of *C. coli* strain CVM29710. We conducted imputation for polymorphic sites with missing frequency ≤10% using BEAGLE [65] as previously reported [66]. A total of 286,393 gapless SNPs (∼17% of the average *C. coli* genome size) were used for recombination analyses. The coancestry of genome-wide haplotype data was inferred based on alignments using chromosome painting and FineStructure [67] as previously described[68]. Briefly, ChromoPainter was used to infer chunks of DNA donated from a list of 33 donor groups normalized for sample size to each of 677 ST-CC-828 and 12 CC-1150 recipient haplotypes. Results were summarized into a coancestry matrix containing the number of recombination-derived chunks from each donor to each recipient individual. FineStructure was then used for 100,000 iterations of both the burn-in and Markov chain Monte Carlo chain to cluster individuals based on the co-ancestry matrix. The results are visualized as a heat map with each cell indicating the proportion of DNA “chunks” (a series of SNPs with the same expected donor) a recipient receives from each donor.

### Analysis of covariation in bacterial genomes

Non-random allele associations can result from selection and clonal population structure. To control for the latter, our approach identified SNP combinations in independent genetic backgrounds by accounting for the sequence variation associated with the inferred phylogeny. Based on the alignment of 677 genomes of *C. coli* CC-828, a first phylogenetic tree was created using PhyML [69]. ClonalFrameML [64] was then applied to correct the tree by accounting for the effect of recombination, and also to infer the ancestral sequence of each node. Covariance was assessed for pairs of biallelic sites across the genome using a Continuous Time Markov Chain (CTMC) model as follows. Briefly, let *A* and *a* denote the two alleles of the first site and *B* and *b* denote the two alleles of the second site, so that there are four states in total for the pair of sites (*ab*, *Ab*, *aB* and *AB*). The four substitution rates from *A* to *a*, from *a* to *A*, from *B* to *b* and from *b* to *B* are not assumed to be identical, to allow for differences in substitution rates in different parts of the genome and also to allow for non-equal rates of forward and backward substitution (for example as a result of recombination opportunities). Assuming no epistatic effect between the two sites (ε=1), the model M_0_ has four free parameters (α_1_, α_2_, β_1_ and β_2_) representing independent substitutions at the two sites. We expand model M_0_ with an additional fifth parameter ε>1 into model M_1_ which is such that the state *AB* where the first site is allele *A* and the second site is allele *B* is favored relative to the other three sites *ab, aB* and *Ab*. Specifically, the state *AB* has a probability increased by a factor ε^2^ in the stationary distribution of the CTMC of model M_1_ compared to model M_0_.

Both models M_0_ (with 4 parameters) and M_1_ (with 5 parameters) are fitted to the data using maximum likelihood techniques, where the likelihood is equal to the product for every branch of the tree of the state at the bottom of the branch given the state at the top. The two fitted models M_0_ and M_1_ are then compared using a likelihood-ratio test (LRT) as follows: since M_0_ is nested with M_1_, two times their difference in log-likelihood is expected to be distributed according to a chi square distribution with number of degrees of freedom equal to the difference in their dimensionality, which is one. This LRT returns a *p-*value for the significance of a covariation effect, and a Bonferroni correction is applied to determine a conservative cutoff of significance that accounts for multiple testing. Furthermore, the test is applied only to pairs of sites separated by >20kb to reduce the chance that they were the result of a single recombination event, consistent with estimates of the length of recombined DNA sequence in quantitative bacterial transformation experiments [39, 70] and evidence from *Campylobacter* genome analyses that show that LD for pairs of sites decreases with distance to approximately 20kbp and then remains at the same level for very distant sites [42]. It is still possible of course that rare recombination events would stretch 20kbp [39–41], but for this to have an effect on the analysis of epistasis it would have to have happened several times for the same pairs of sites against different genomic background which becomes quite unlikely just by chance. This phylogenetically aware approach to testing for covariance presents the advantage to naturally account for both population structure and the effect of recombination [71]. The script implementing this coevolution test is available in R at: https://github.com/xavierdidelot/campy.

### Quantifying covariation between recombined and unrecombined genomic regions

The results of the introgression and covariation analyses were combined so that for each pair of significantly covarying SNPs (*p-*value <10^-8^), haplotype frequency was calculated among the 689 recipient introgressed *C. coli* clade-1 strains as well as among the donor *C. coli* (ancestral) and *C. jejuni* strains, respectively. If the most frequent haplotype of the pair is the same between the donor *C. coli* (ancestral) and *C. jejuni*, it was classified as ‘no polymorphism’. Otherwise, if the most frequent haplotype accounted for >90% among the recipients, it was classified as either ‘*C. jejuni* (>90%)’ or ‘*C. coli* (>90%)’ if it was the same as that of donor *C. jejuni* or *C. coli* (ancestral) (inset in Figure 2C). If the most frequent haplotype accounted for ≤90% among the recipients, the top two most frequent haplotypes (written as major and minor haplotype in this manuscript) were indicated as either “*C. jejuni* / *C. coli*”, “*C. jejuni* / other”, “*C. coli* / *C. jejuni*”, “*C. coli* / other”, “other / *C. coli*”, “other / *C. jejuni*”, and “other / other”, and the frequency of the major and minor haplotypes were calculated. For example, where the haplotype frequencies were as follows, AA=285, TA=192, TG=181, AG=27, A-=2, --=1, -A=1, AA is the major haplotype, frequency of which is 41.3%

### Mutagenesis and complementation cloning

Genes *cj1167*, *cj1168c* (here designated *selF* for <u>sel</u>enium transport for formate dehydrogenase), *cj1171c* (*ppi*), *cj1507c* (*modE*), *cj1508c* (*fdhD*) and *cj1500* (*fdhT*) were deleted by allelic exchange mutagenesis, with the majority of the open reading frame replaced by an antibiotic resistance cassette. Mutagenesis plasmids were generated by the isothermal assembly method using the HiFi system (NEB, UK). In brief, flanking regions of target genes were PCR amplified from genomic DNA using primers with adaptors homologous to either the backbone vector pGEM3ZF or the antibiotic resistance cassette (Data S5). pGEM3ZF was linearised by digestion with HincII. The kanamycin and chloramphenicol resistance cassettes were PCR amplified from pJMK30 and pAV35, respectively [72]. Four fragments consisting of linearised pGEM3ZF, antibiotic resistance cassette and 2 flanking regions were combined in equimolar amounts and mixed with 2 x HiFi reagent (NEB, UK) and incubated at 50°C for 1 hour. The fragments combine such that the gene fragments flank the antibiotic resistance cassette, in the same transcriptional orientation, within the vector. Mutagenesis plasmids were transformed into *C. jejuni* NCTC 11168 by electroporation. Spontaneous double-crossover recombinants were selected for using the appropriate antibiotic and correct insertion into the target gene confirmed by PCR screening. For genetic complementation of mutants, genes *cj1168c* (*selF*), *cj1507c* (*modE*), *cj1508c* (*fdhD*) and *cj1500* (*fdhT*) were PCR amplified from genomic DNA, restriction digested with MfeI and XbaI, then ligated into similarly digested pRRA [73] (Data S5). The orientation of insertion allowed the target gene to be expressed constitutively by a chloramphenicol resistance gene-derived promoter within the vector. Complementation plasmids were transformed into *C. jejuni* by electroporation. Spontaneous double-crossover recombinants were selected for using apramycin and correct insertion into the ribosomal locus confirmed by PCR screening.

### Growth of *C. jejuni*

Microaerobic growth cabinets (Don Whitley, UK) were maintained at 42°C with an atmosphere of 10% O_2_, 5% CO_2_ and 85% N_2_ (v/v). *C. jejuni* was grown on Columbia-base agar containing 5% v/v defibrinated horse blood. Selective antibiotics were added to plates as appropriate at the following concentrations: 50 µg ml^-1^ kanamycin, 20 µg ml^-1^ chloramphenicol, 60 µg ml^-1^ apramycin. Muller-Hinton (MH) broth supplemented with 20 mM L-serine was used as a rich medium. Minimal medium was prepared from a supplied MEM base (51200-38, Thermo Scientific, UK) with the following additions: 20 mM L-serine, 0.5 mM sodium pyruvate, 50 µM sodium metabisulfite, 4 mM L-cysteine. HCl, 2 mM L-methionine, 5 mM L-glutamine, 50 µM ferrous sulfate, 100 µM ascorbic acid, 1 µM vitamin B12, 5 µM sodium molybdate, 1 µM sodium tungstate. Selenium was then added as appropriate from stocks of sodium selenate or sodium selenite prepared in dH_2_O. For assays, cells were washed and suspended in sterile phosphate buffered saline (PBS, Sigma-Aldrich).

### Respiration rates with formate

Cells were first grown in MH broth for 12 hours, then washed thoroughly in PBS before inoculating minimal media without an added selenium source. The appropriate concentrations were determined by serial dilution trials, and it was subsequently found that *C. jejuni* has a strong preference for selenite over selenate, as equivalent FDH activity requires some 1000-fold greater concentration of selenate than selenite (Figure S10). These cultures were grown for 8 hours before the cells were thoroughly washed again, then used to inoculate further minimal media, with a selenium source added as appropriate, and grown for 10 hours. This passaging was necessary to remove all traces of selenium from the inoculum, such that control cultures without selenium added had negligible (FDH) activity. Assay cultures were again thoroughly washed before the equivalent of 20 ml at an optical density of 0.8 at 600 nm was finally suspended in 1 ml of PBS. Formate dependent oxygen consumption by whole cells was measured in a Clark-type electrode using 20 mM sodium formate as electron donor. The electrode was calibrated with air-saturated PBS assuming 220 nmol dissolved O_2_ ml^−1^ at 37°C. In the electrode, 200 µl of the dense cell suspension was added to 800 µl air-saturated PBS for a final volume of 1 ml. The chamber was sealed and the suspension allowed to equilibrate for 2 minutes. The assay was initiated by the addition of 20 µl of 1 M sodium formate (prepared in PBS) and the rate of oxygen consumption recorded for 90 s. The total protein concentration of the cell suspensions was determined by Lowry assay and the specific rate of formate-dependent oxygen consumption expressed as nM oxygen consumed min^−1^ mg^−1^ total protein.

## Acknowledgments

**Funding:** Wellcome Trust grant 088786/C/09/Z (SS), Medical Research Council (MRC) grant MR/M501608/1 (SS), Medical Research Council (MRC) grant MR/L015080/1 (SS), Food Standards Agency project FS101087 (SS), Biotechnology & Biological Sciences Research Council (BBSRC) grant BB/S014497/1 (DK).

## Author contributions

Conceptualization and study design: SS, XD, DF, KY, DK, AT Sample collection: SS, AvV, NW

Laboratory work: AT, LM, MH, BP Data archiving: BP, KJ

Data analysis: AT, XD, KY, LM, JKC, EM, SP, SB, SS, JC

Data interpretation: KY, XD, SS, MM, JP, DK, AT, DF Writing: SS, CK, AT, DK

Competing Interest Statement: The authors declare that they have no competing interests.

## Data and materials availability

Short-read sequence data for all isolates sequenced in this study are deposited in the sequence read archive (SRA) and can be found associated with NCBI BioProjects PRJNA689604 (https://www.ncbi.nlm.nih.gov/bioproject/PRJNA689604) and PRJEB11972 (https://www.ncbi.nlm.nih.gov/bioproject/PRJEB11972). These were augmented with 498 previously published genomes and assembled genomes are available on Figureshare (doi.org/10.6084/m9.Figureshare.13521683). Accession numbers for all genomes are included in Data S4. Source data are provided for this paper (Data S2).

## Notes

https://www.ncbi.nlm.nih.gov/bioproject/PRJNA689604

https://www.ncbi.nlm.nih.gov/bioproject/PRJEB11972

### Competing Interest Statement

The authors have declared no competing interest.

### Summary of Updates

Revised as part of the peer review process and additional authors added to reflect the work conducted.

