## Supplementary material for "A two-hit epistasis model prevents core genome disharmony in recombining bacteria"

**This file includes:**

Figs. S1 to S9

Data S5

**Other Supplementary Materials for this manuscript include the following:**

Data S1 to S4

<insert page break then Fig. S1 here>

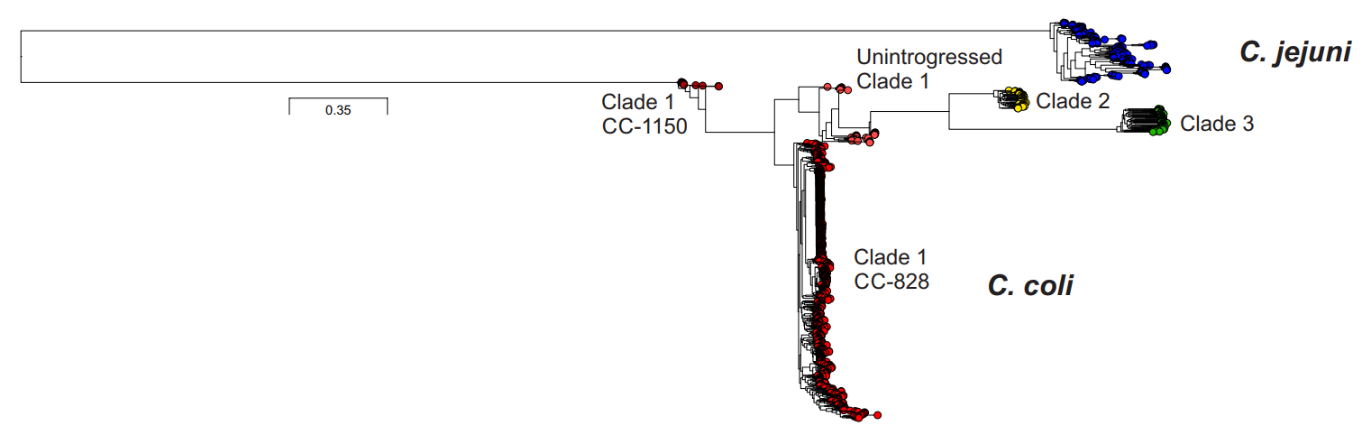

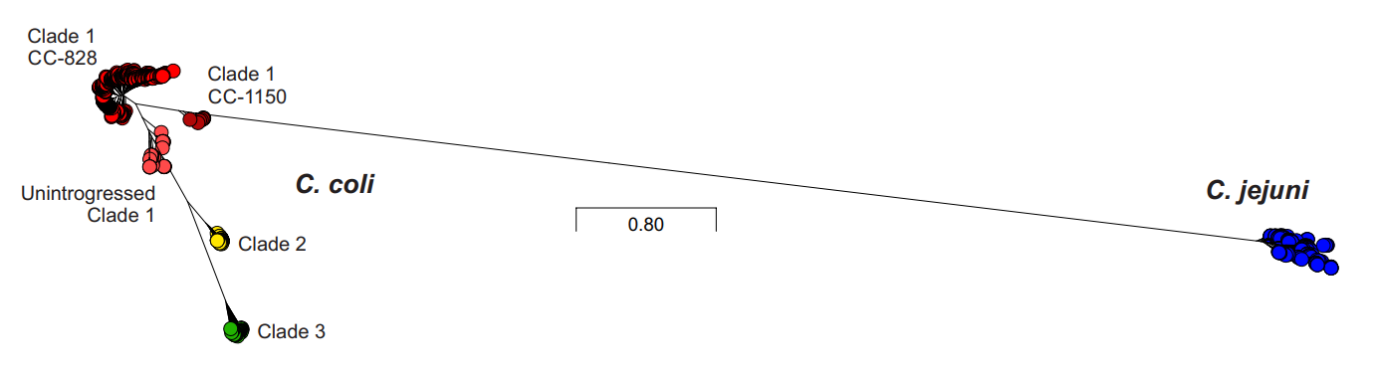

Fig. S1.

Type or paste caption here. Create a page break and paste in the Fig above the caption.

<insert page break then Fig. S2 here>

Fig. S2.

Type or paste caption here. Create a page break and paste in the Fig above the caption.

<insert page break then Fig. S3 here>

Fig. S3.

Type or paste caption here. Create a page break and paste in the Fig above the caption.

**Fig S1. RAxML phylogeny of 973 *C. jejuni* and *C. coli* clades.** The phylogenetic tree was constructed using an approximation of the maximum-likelihood (ML) method implemented in RAxML using the GTRGAMMA model, from an imputed whole-genome alignment of 973 genomes to the *C. coli* CVM N29710 reference genome (ST-828 clonal complex) and a total of 286,393 variable sites. Three clades of *C. coli* are included: 689 isolates from introgressed clade 1 (677 from ST-828 complex and 12 from ST-1150 complex) and 35 isolates from unintrogressed clade 1 (red), 45 isolates from clade 2 (yellow) and 58 isolates from clade 3 (green). A selection of 146 *C. jejuni* from a representation of 30 different clonal complexes is included (blue). The scale represents the number of substitutions per site.

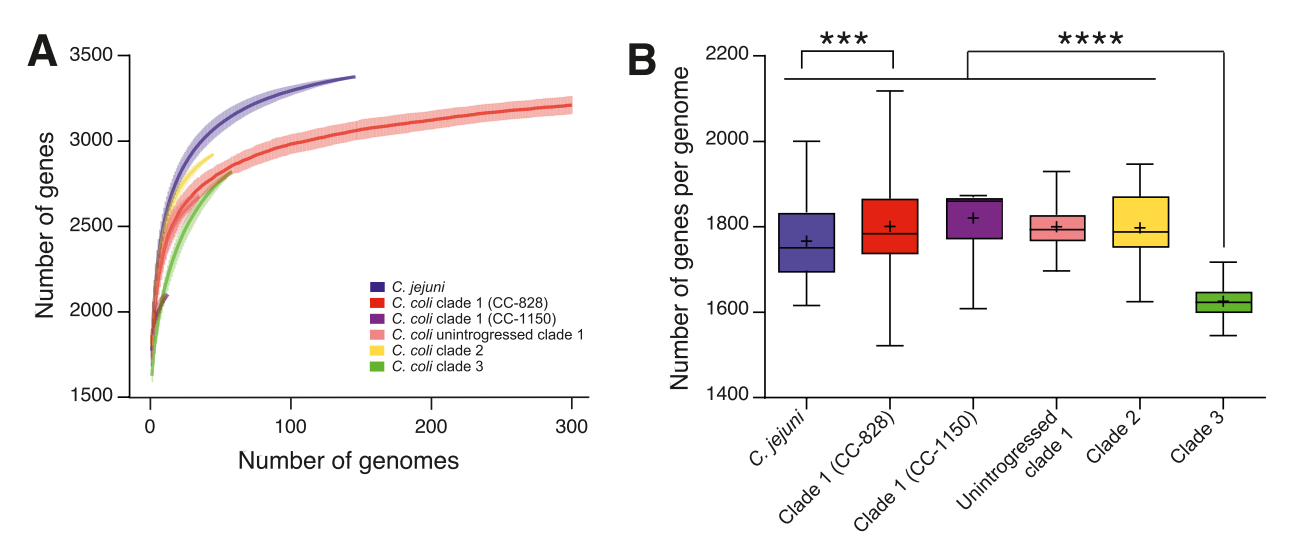

**Fig S2. Population genomics of *C. jejuni* and *C. coli.*** (A) Accumulation curves of the clade-specific pan-genome content of *Campylobacter* lineages, using the same colour code as panel A. Randomized genome sampling was used to obtain the average number of genes for each comparison (plain lines) and standard deviations (dotted lines). (B) The average number of genes/genome, identified by BLAST, is shown for *C. jejuni* and the different *C. coli* clades. Asterisks denote significant difference between distributions as inferred from a Dunn’s multiple comparisons test after a Kruskal-Wallis test, as follows: ***: p<0.001; ****: p<0.0001.

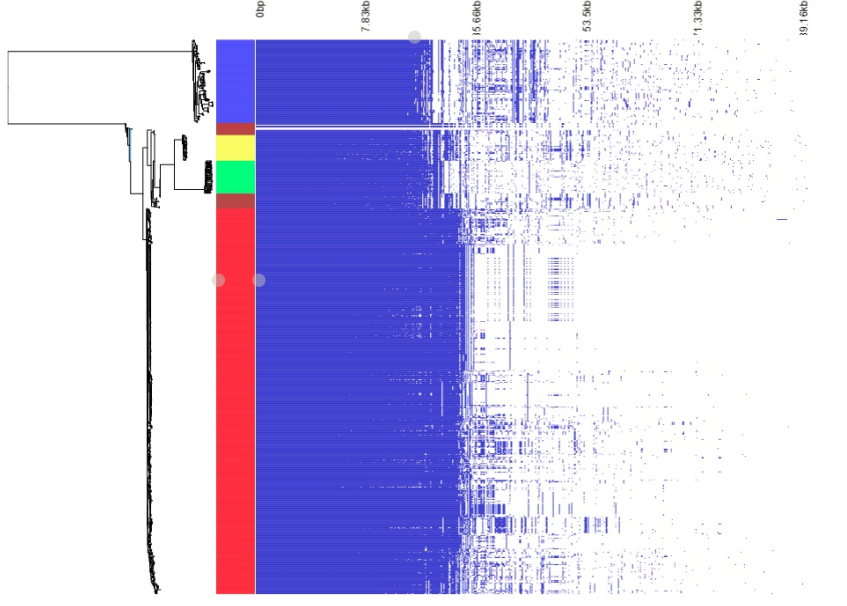

**Fig S3. Pan genome variation among *C. jejuni* and *C. coli* isolates**

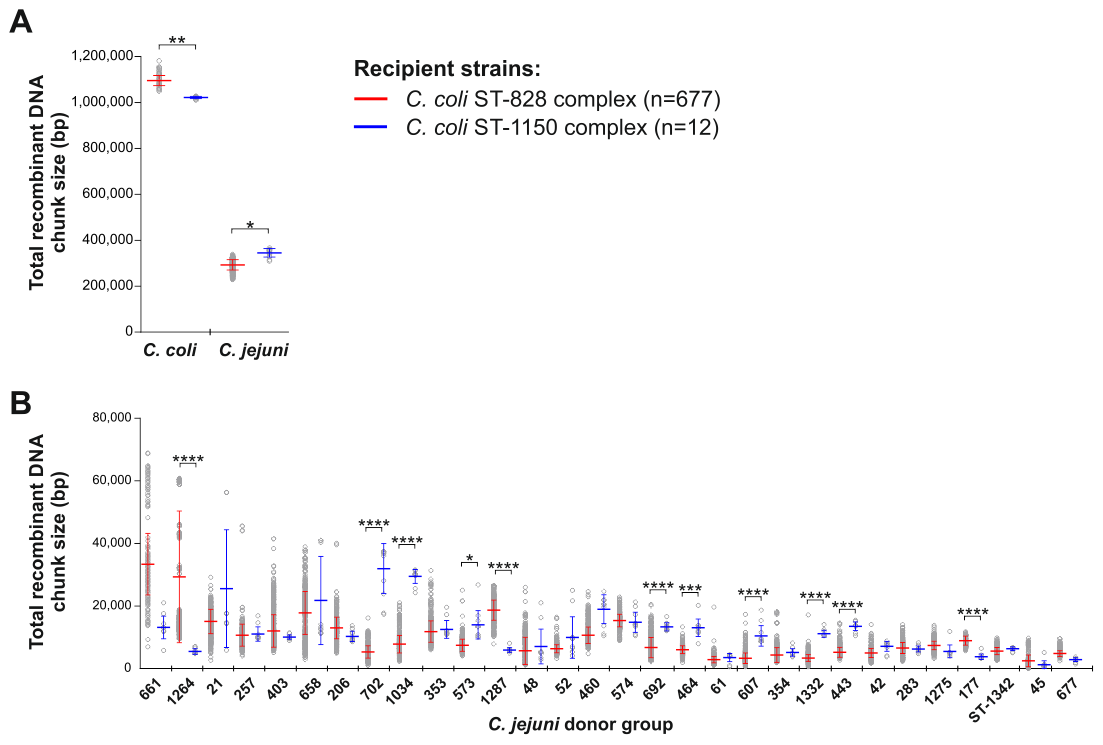

**Fig S4. Chromosome painting inference of recombination-derived *C. jejuni* sequence in recipient *C. coli* genomes.** (A) Length of DNA chunks recombining from 30 *C. jejuni* donor groups into *C. coli* ST-828 and ST-1150 clonal complexes and (B) the relative importance of different *C. jejuni* clonal complexes as donors.

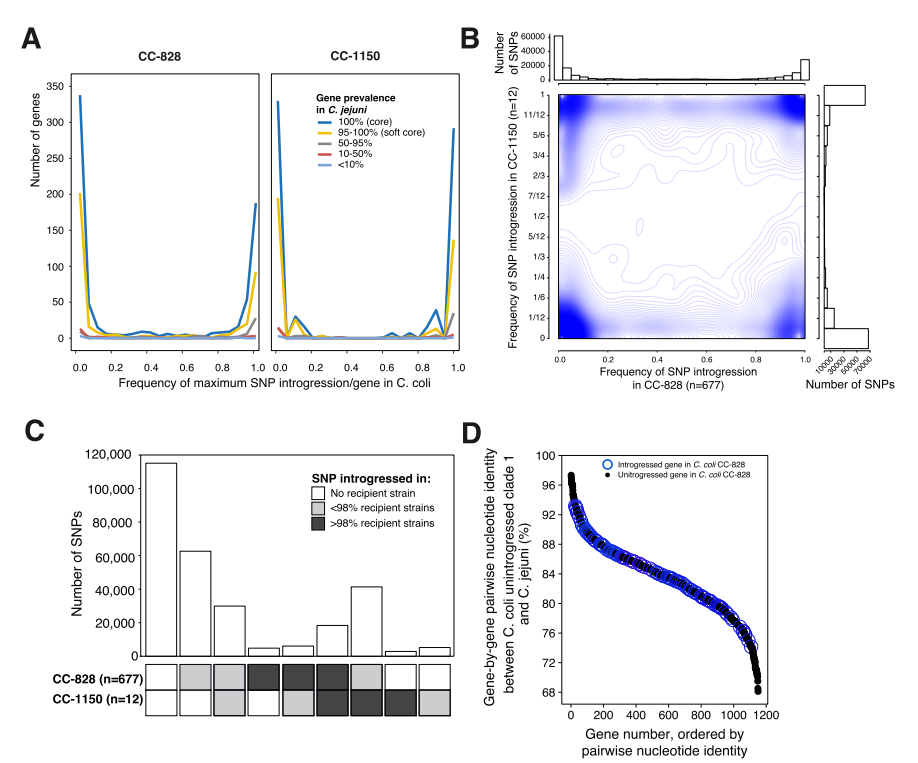

**Fig S5. Genome-wide introgression from *C. jejuni* to *C. coli*.** (A) The number of genes with different frequencies of maximum SNP introgression/gene in *C. coli* as a function of gene frequency in *C. jejuni*. Highly introgressed genes in CC-828 and CC-1150 tend to be core in *C. jejuni*. (B) Density plot (n=1000 bins) of specific and shared introgression events in CC-828 (x-axis) and CC-1150 (y-axis). The frequency of SNP introgression/gene is shown for both lineages. Close blue lines denote a high density of points. (C) Shared introgression between *C. coli* CC-828 and CC-1150. The number of SNPs being shared between the two lineages at various frequencies is shown in y-axis. (D) Pairwise nucleotide identity between *C. jejuni* and ancestral (unintrogressed) clade 1 *C. coli* core genes (black circles). Genes found to be introgressed in clade 1 CC-828 are highlighted in blue.

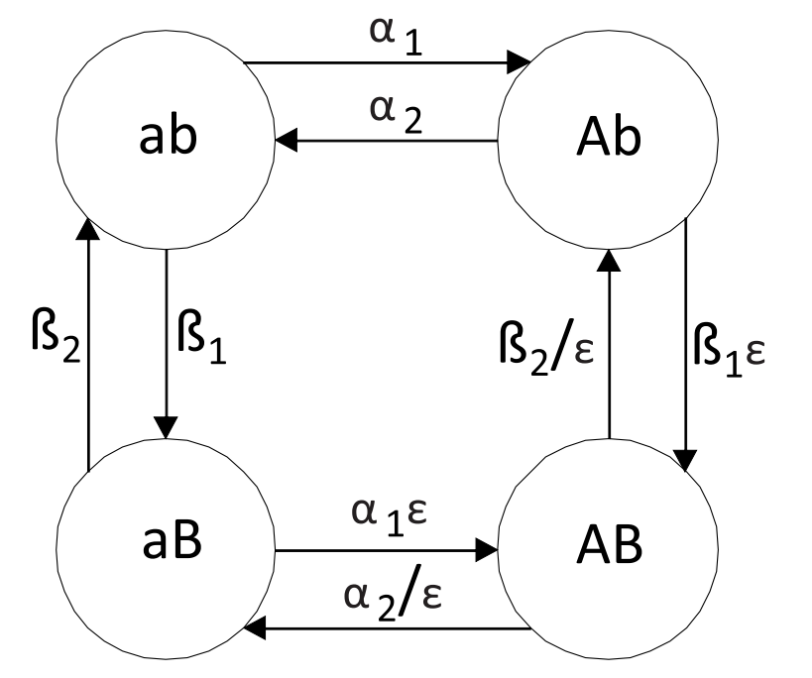

**Fig S6.** **Schematic of the continuous time Markov chain (CTMC) model used to analyse covariation in the *Campylobacter* genomes.** Covariation was assessed for pairs of biallelic sites separated by at least 20kb along the genome while accounting for the effect of population structure and recombination.

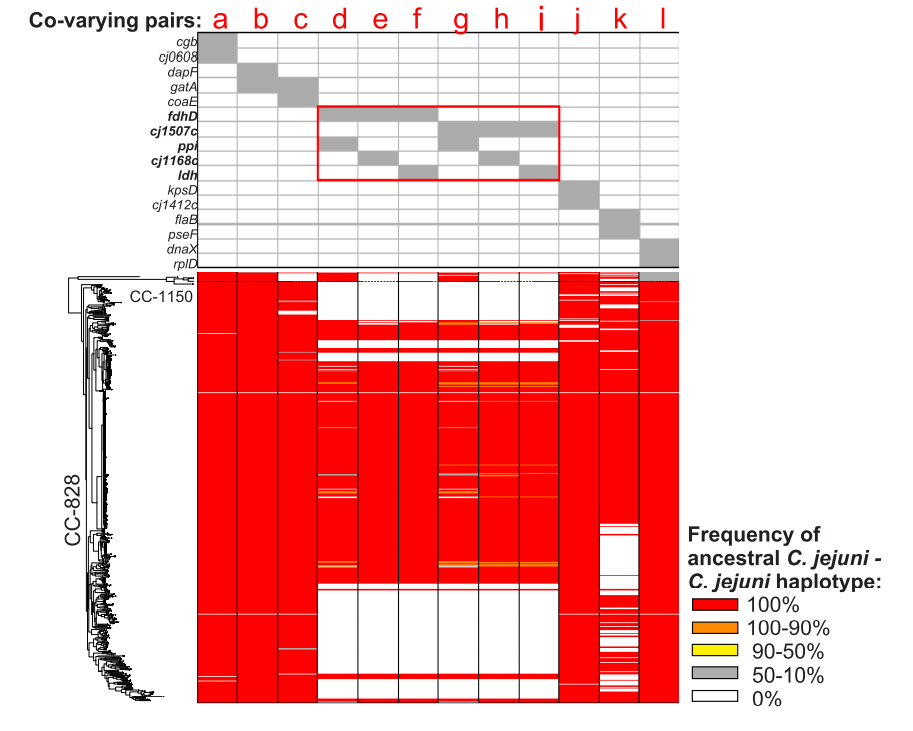

**Fig S7. Putative epistatic SNP pairs in introgressed *C. coli* genomes where the major haplotype was *C. jejuni* indicating that both co-varying sites had *C. jejuni* ancestry*.*** 16 genes pairs in which the *C. jejuni* - *C. jejuni* haplotype is dominant (a to l), and frequency of ancestral *C. jejuni - C. jejuni* haplotype among the epistatic SNP pairs in co-varying genes is shown as the 5 categories for each *C. coli* strain mapped to the phylogeny (ML tree). The red indicates 100% of the epistatic SNPs in the co-varying gene are ancestral *C. jejuni - C. jejuni* haplotype.

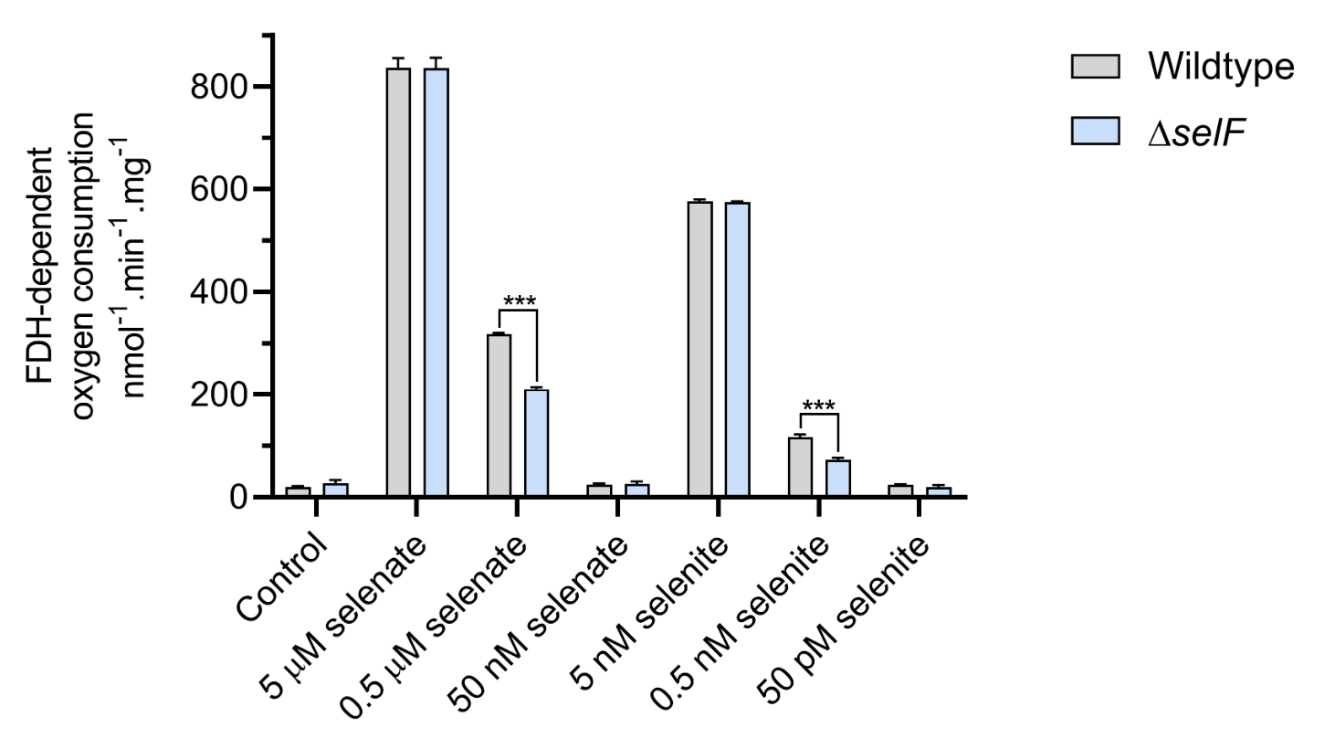

**Fig S8. Effect of selenite or selenate concentration on the FDH activity of the *selF* mutant and parental wildtype grown in minimal media.**

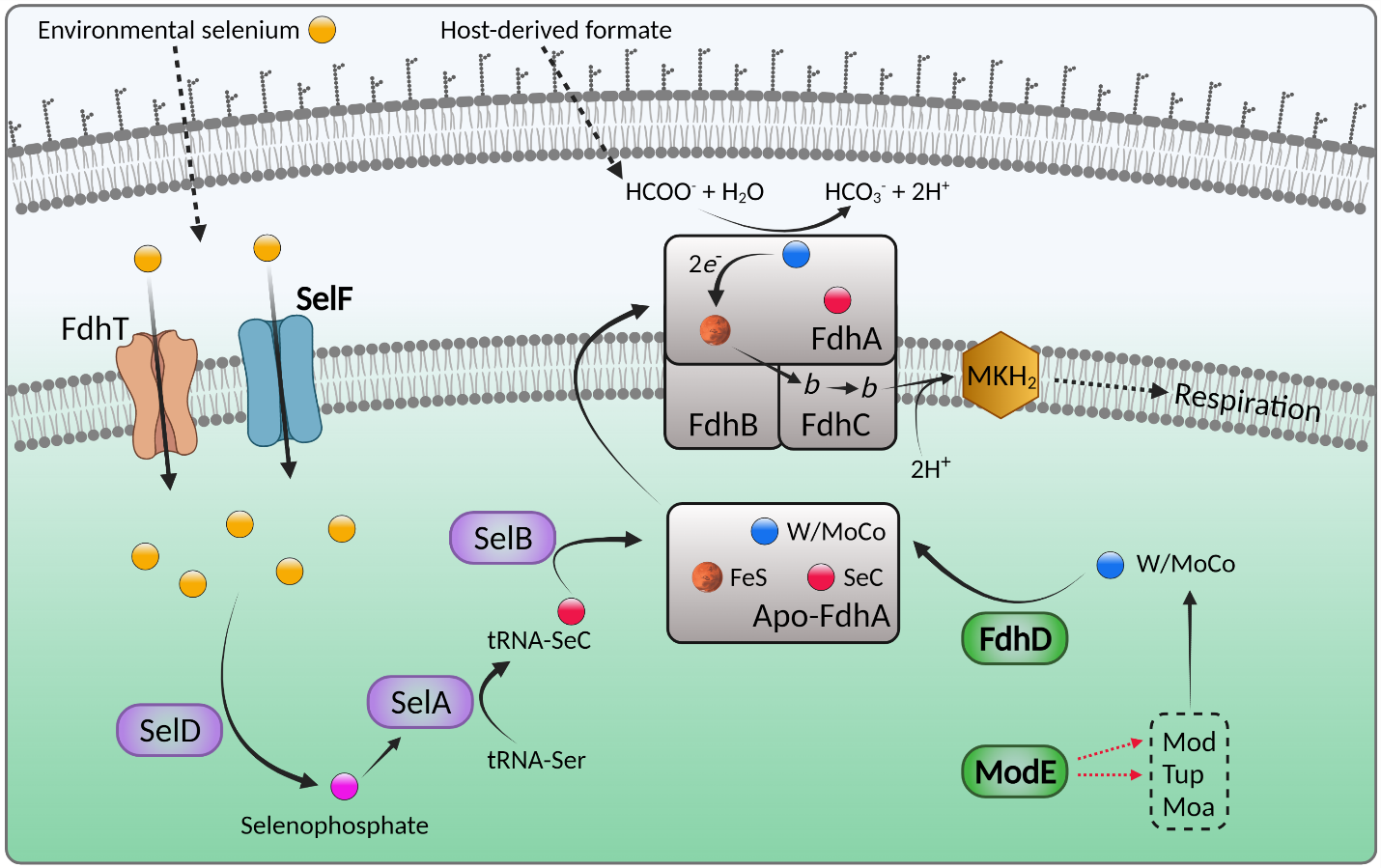

**Fig S9. Model for epistatically linked genes involving FDH biogenesis and activity.** Host derived formate is converted to bicarbonate in the periplasm by the FDH complex to release electrons which are transferred from the iron sulfur (FeS) cluster of FdhA to the *b*-type hemes of FdhC and into the menaquinone (MK) pool where they can ultimately be used to reduce molecular oxygen via terminal oxidases. FdhA contains a selenocysteine residue and Mo/W-pterin cofactor (W/MoCo) at its active site, both of which are essential for catalysis. ModE is a DNA binding regulator which represses the Mo and W transporters Mod and Tup to regulate the cellular pool of Mo/W. W/MoCo is generated by the Moa pathway and inserted into apo-FdhA by the sulphur-transferase FdhD. Environmental selenite (the most abundant oxyanion) or selenate diffuses into the periplasm where it must be actively imported to the cytoplasm by FdhT and SelF. Putatively, FdhT is low affinity and functions efficiently when ample selenium is available. When selenium is limited, SelF can import sufficient selenium to maintain FDH production and activity. Cytoplasmic selenium is converted to selenophosphate by SelD, which is used by SelA to generate tRNA-SeC from tRNA-Ser. During translation of FdhA, tRNA-SeC is incorporated by the specific elongation factor SelB. Apo-FdhA with W/MoCo and SeC inserted is then transported to the periplasm by the TAT translocation system and incorporated into the FDH complex.

| **Mutagenesis** | **Sequence 5' - 3'** |
| --- | --- |
| 1167F1 | GAGCTCGGTACCCGGGGATCCTCTAGAGTCgaatttgccatagaaacaa |
| 1167R1 | AAGCTGTCAAACATGAGAACCAAGGAGAATgttctgaacaaattccttg |
| 1167F2 | GAATTGTTTTAGTACCTAGCCAAGGTGTGCgctatcataggaaaagatgg |
| 1167R2 | AGAATACTCAAGCTTGCATGCCTGCAGGTCagtgggactttaggttctg |
| 1168F1 | GAGCTCGGTACCCGGGGATCCTCTAGAGTCtgatagatatcaattcttacgc |
| 1168R1 | AAGCTGTCAAACATGAGAACCAAGGAGAATcaactctctaaggtcattaaga |
| 1168F2 | GAATTGTTTTAGTACCTAGCCAAGGTGTGCggacaaaatgaagagttga |
| 1168R2 | AGAATACTCAAGCTTGCATGCCTGCAGGTCgaacatggagattctcagtt |
| 1171F1 | GAGCTCGGTACCCGGGGATCCTCTAGAGTCgaaaagatagcgcagatatta |
| 1171R1 | AAGCTGTCAAACATGAGAACCAAGGAGAATgtattcgtttactttgcttgt |
| 1171F2 | GAATTGTTTTAGTACCTAGCCAAGGTGTGCttagacataaaaatttgcgat |
| 1171R2 | AGAATACTCAAGCTTGCATGCCTGCAGGTCttgttattagcactataatcttgtc |
| 1507F1 | GAGCTCGGTACCCGGGGATCCTCTAGAGTCcttaatgctgaaggtgtagtt |
| 1507R1 | AAGCTGTCAAACATGAGAACCAAGGAGAATacaattctctcatatatgagatg |
| 1507F2 | GAATTGTTTTAGTACCTAGCCAAGGTGTGCttgatggaaaattactctatttt |
| 1507R2 | AGAATACTCAAGCTTGCATGCCTGCAGGTCtttcttcatcagcgtaaag |
| 1508F1 | GAGCTCGGTACCCGGGGATCCTCTAGAGTCggttatttaagtaaaatcaaacg |
| 1508R1 | AAGCTGTCAAACATGAGAACCAAGGAGAATttgagttgtaaataacggatc |
| 1508F2 | GAATTGTTTTAGTACCTAGCCAAGGTGTGCgatcaatatttatagtggagaattta |
| 1508R2 | AGAATACTCAAGCTTGCATGCCTGCAGGTCgaagcatgtaataaactacctgt |
| 1500F1 | GAGCTCGGTACCCGGGGATCCTCTAGAGTCcgttctatgattgcacttg |
| 1500R1 | AAGCTGTCAAACATGAGAACCAAGGAGAATcttgtttaacggcaagttt |
| 1500F2 | GAATTGTTTTAGTACCTAGCCAAGGTGTGCggtcatactagcaaactcataga |
| 1500R2 | AGAATACTCAAGCTTGCATGCCTGCAGGTCggcagcttatgaaaagatg |
| **Complementation** | **Sequence 5' - 3'** |
| 1168fwd | *acac*TCTAGAttaaggaagacaATGCAAGAAATCATTTCTTT |
| 1168rev | *acac*CAATTGgctatcataggaaaagatgg |
| 1507fwd | *acac*TCTAGAttaaggaagacaATGGATAAAAATCAAGAAAAAG |
| 1507rev | *acac*CAATTGcctatatttaaagatttaaacatcttat |
| 1508fwd | *acac*TCTAGAttaaaggataaggtatggatcc |
| 1508rev | *acac*CAATTGcacaatctaatttttcattgc |
| 1500fwd | *acac*TCTAGAttaaggaagacaTTGAATTCTTTTAAACAAAAATATTTAATC |
| 1500rev | *acac*CAATTGggacaagcttcaccttgtaa |

Data S5. Primers used in this study.

Data S1. (separate file) Maximum SNP frequency and covarying p-value per gene.

Data S2. (separate file) Source data.

Data S3. (separate file) Genes with covarying SNPs identified in this study.

Data S4. (separate file) Isolates and genomes used in this study.
